# Design, optimization, and analysis of large DNA and RNA nanostructures through interactive visualization, editing, and molecular simulation

**DOI:** 10.1101/2020.01.24.917419

**Authors:** Erik Poppleton, Joakim Bohlin, Michael Matthies, Shuchi Sharma, Fei Zhang, Petr Šulc

## Abstract

This work seeks to remedy two deficiencies in the current nucleic acid nanotechnology software environment: the lack of both a fast and user-friendly visualization tool and a standard for common structural analyses of simulated systems. We introduce here oxView, a web browser-based visualizer that can load structures with over 1 million nucleotides, create videos from simulation trajectories, and allow users to perform basic edits to DNA and RNA designs. We additionally introduce open-source software tools for extracting common structural parameters to characterize large DNA/RNA nanostructures simulated using the coarse-grained modeling tool, oxDNA, which has grown in popularity in recent years and is frequently used to prototype new nucleic acid nanostructural designs, model biophysics of DNA/RNA processes, and rationalize experimental results. The newly introduced software tools facilitate the computational characterization of DNA/RNA designs by providing multiple analysis scripts, including mean structures and structure flexibility characterization, hydrogen bond fraying, and interduplex angles. The output of these tools can be loaded into oxView, allowing users to interact with the simulated structure in a 3D graphical environment and modify the structures to achieve the required properties. We demonstrate these newly developed tools by applying them to *in silico* design, optimization and analysis of a range of DNA and RNA nanostructures.

## I. INTRODUCTION

The field of nucleic nanotechnology^1^ uses DNA and RNA as building blocks to construct nanoscale structures and devices. Using the high programmability of pairing combinations between oligonucleotides, it is possible to construct 2D and 3D nanostructures up to several thousand nucleotides. Over the past three decades, designs of increasing complexity have been proposed, such as DNA/RNA tiles and arrays^2^, DNA multi-bundle origamis^3^, wireframe nanostructures^4,5^ single-stranded tile (SST) nanostructures^6^, single-stranded DNA (ssDNA) and RNA (ssRNA) origami structures^7^, and larger multi-origami tile assemblies^8^. The nanostructures have promising applications ranging from photonic devices^9^ to drug delivery^10^.

There are many available nucleic acid nanotechnology design tools, including CaDNAno^11^, Tiamat^12^, vHelix^13,14^, Adenita^15^, MagicDNA^16^, and the CAD converters DAEDALUS^17^ and PERDIX^18^. CaDNAno is frequently used to design very large structures on either a square or hexagonal lattice, which requires components be made of parallel helices. Tiamat is an intuitive lattice-free design tool that supports both DNA and RNA. MagicDNA is a Matlab-based tool that specializes in the design of large 3D structural components on a 3D cubic lattice using CaDNAno-like parallel DNA bundles as the base unit of each edge. VHelix and Adenita are DNA design plugins for the commercial design platforms Maya and SAMSON. VHelix facilitates conversion of polyedral meshes to DNA sequences, with further free-form editing available in Maya. Adenita combines the functionality of CAD converters with free-form design, allowing users to load structures from a variety of sources with additional editing tools available in the SAMSON interface. DAEDALUS and PERDIX are software that facilitate conversion of meshes designed in CAD software into DNA representations. Currently, the nanotechnology field lacks a universal method for assembling structures made in different design tools, especially if small changes need to be made. Continued development of tools is thus necessary to integrate the previous efforts and enable design of more complex DNA and RNA nanostructures. Additionally, with the exception of Tiamat, all available tools focus only on DNA nanostructure designs.

Molecular simulations have proved indispensable in the field of nucleic acid nanotechnology, providing detailed information about bulk structural characteristics^19,20^, folding pathway kinetics^21,22^, conformational space and kinetics of complex nanostructures^23–25^, and active devices such as DNA walkers^26,27^. Due to the size of the designed nanostructures and the laboratory timescales involved, traditional fully atomistic simulation methods are often infeasible for nucleic acid nanotechnology applications. To remedy this, several coarsegrained models have been developed^28–36^, each of which with a unique focus on a specific part of the DNA nanostructural design and characterization pipeline. In particular, the oxDNA/oxRNA models have grown in popularity in recent years and have been used for studying DNA/RNA nanostructures and devices^23,32,37–39^ as well as RNA/DNA biophysics^30,40,41^. The models represent each nucleotide as a single rigid body, where the interactions between nucleotides are empirically parameterized to reproduce basic structural, mechanical and thermodynamic properties of DNA and RNA (Fig. 1).

**FIG. 1.**
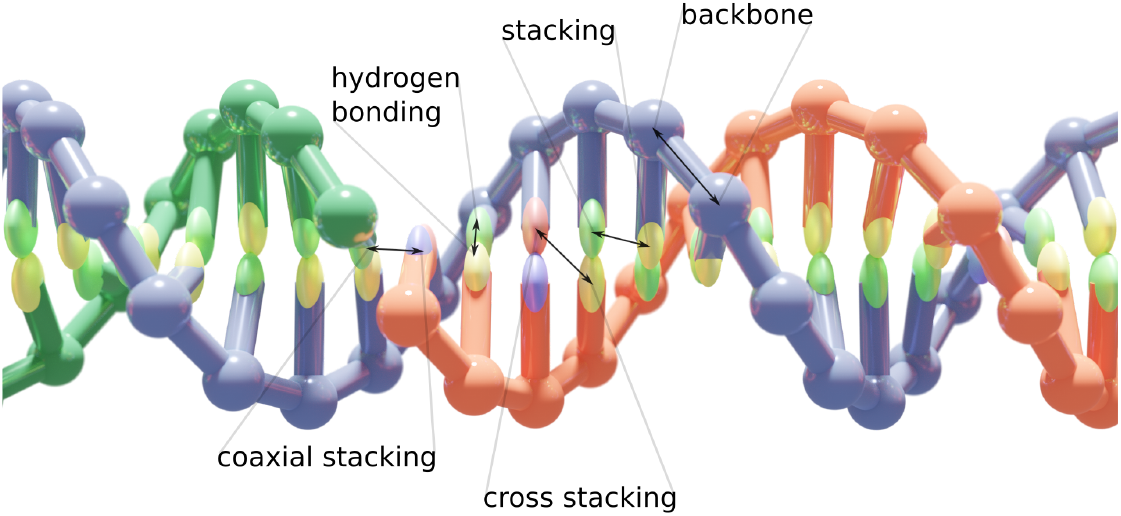
The oxDNA model. A DNA duplex as modeled in oxDNA with labels corresponding to the coarse-grain potentials defining the force field. OxDNA is a “two-bead” model with each nucleotide represented as two beads with specific interaction sites that approximate the geometry and interactions of the 20+ atoms that make up each nucleotide. The coarse-grained force field is parameterized to reconstruct the structural and dynamic properties of both single- and double-stranded DNA and RNA.

However, the standalone simulation package only provides simulation trajectory with recorded 3D positions of all nucleotides in the simulation. Users usually have to develop in-house evaluation tools that post-process the simulation trajectory to extract desired properties of the studied nanostructures.

In this paper, we present two open-source tools to fill these unmet needs in the field of DNA/RNA nanotechnology and illustrate their use for design and optimization of DNA and RNA nanostructures. The first tool we introduce here is oxView, a browser-based visualization and editing platform for DNA and RNA structural design and analysis of nanostructures simulated in oxDNA/oxRNA. The tool is able to accommodate nanostructures containing over a million nucleotides, which is beyond the reach of most other visualization tools. It allows the user to load multiple large nanostructures simultaneously and edit them by addition or deletion of individual nucleotides or entire regions, providing a way to create new, more complex designs from smaller, individually designed subunits, even from different design tools. All of the previously mentioned design tools can be converted to the oxDNA format using either built-in tools (Adenita, MagicDNA, vHelix), the TacoxDNA webserver^42^ (CaDNAno, Tiamat, vHelix), or by converting first to PDB using built-in tools and then to oxDNA using TacoxDNA (DAEDALUS, PERDIX). The visualization tool is integrated with oxDNA/oxRNA simulations and loads long simulation trajectories quickly (including files which are tens of gigabytes in size) for interactive analysis and video export of nanostructure dynamics. It can also load data overlays from the analysis scripts introduced in this paper, allowing users to interactively explore features such as hydrogen bond occupancy and structure flexibility and then use this information to iteratively redesign nanostructures based on simulation feedback using oxView. Finally, oxView implements rigid-body dynamics code so that individual parts of the structures can be selected and interactively rearranged. The structure will then be relaxed on-the-fly using rigid-body dynamics to a conformation which can be used as an initial structure in simulations.

The second tool introduced here is a set of standardized structure-agnostic geometry analysis scripts for oxDNA/RNA which cover a number of common molecular simulation use cases. Many groups that work with oxDNA/RNA have developed their own analysis tools in-house, resulting in many duplicate functionalities and scripts that are limited to single experiments. To facilitate the simulation-guided design of DNA/RNA nanostructures and lower the barrier of entry into the simulation field, we have developed a toolkit that is easy to use, generically applicable to numerous studied systems, and extensible. The tool set includes the following: 1) calculation of mean structure and root-mean squared fluctuations to quantify structure flexibility; 2) hydrogen-bond occupancy to quantify fraying and bond breaking during the simulation; 3) angle and distance measurements between respective duplex regions in a nanostructure; 4) a covariance-matrix based principle component analysis tool for identification of nanostructure motion modes; and 5) unsupervised clustering of sampled configurations based on structural order parameters or global difference metrics.

We demonstrate the versatility of the analysis tools and visualization platform functionality by analyzing simulations of previously published structure and a few novel designs. In particular, we study two RNA tiles, a Holliday junction, the tethered multi-fluorophore structure, two wireframe DNA origamis, and a single-stranded RNA origami nanostructure. We make no custom modifications to the analysis tools for each of the designs to demonstrate their versatility and general utility for distinct nanostructures. The visualization and analysis software developed in this work is freely available under a public license.

## II. MATERIALS AND METHODS

### A. System and software requirements

The analysis tools were written and tested using the following dependencies: Python 3.7 (minimum version 3.6), NumPy 1.16^43^, MatPlotLib 3.0.3 (minimum version 3.0)^44^, BioPython 1.73^45^, SciKitLearn 0.21.2^46^Pathos 0.2.3^47^, oxDNA 6985 (minimum version June 2019)^31,32,48^.

OxView will run as-is on any modern web browser with We-bGL support; though, we note that Google Chrome performs best at very large structure sizes. To make modifications to the code, the following dependencies are required: JavaScript ES6, Typescript 2.9.0

### B. OxDNA input and output files format

As the analyses discussed here process the data from oxDNA files, it warrants a brief description of the file types encountered when running and analyzing oxDNA files. This will mitigate confusion as to why certain files are used in each case. The three most important files are the input parameter file, the topology file and the trajectory file. The input file is a text file with the simulation’s parameter names and values separated by equal signs. This file defines simulation and physical parameters such as temperature, the force field used, and file I/O information. Input files will be used by any analysis that calculates energy (primarily those concerned with hydrogen bonds) or is optimized by writing its most computationally-intensive operations distributed and compiled with the oxDNA code. Topology files define the nucleotides present in the simulation. It is a text file with the extension .top and contains a header line with the number of nucleotides and the number of strands separated by a space. Following the header, each line defines a nucleotide with four values: the strand number, the base identity, 3’ covalent connection, and 5’ covalent connection (note that oxDNA numbers bases 3’-5’, the reverse of the biochemistry convention). Strand ends have a value of −1. The trajectory file contains the position, orientation and velocity of every nucleotide at each timepoint, based on the interval specified in the input file. Each configuration in the trajectory begins with a three-line header, defining the temperature, simulation box dimensions, and kinetic, potential, and total energies of the system. Each subsequent line defines one nucleotide with 15 parameters: position, orientation defined by two orthogonal vectors, translational velocity and rotational velocity. All parameters are in XYZ coordinates. The TacoxDNA webserver^42^ has a variety of conversion tools from popular nanotechnology design tools and simulation formats into the oxDNA format. Once converted, these files can be visualized and edited using oxView or simulated using oxDNA/oxRNA. However, for large and/or long simulations, these files can become quite unwieldy, requiring tens of gigabytes of storage and therefore are impossible to open and read in a single reading frame. Therefore, all analyses and visualizations described here read trajectory files in a stream, allowing reading of files that would not otherwise fit in the computer’s RAM.

### C. Example Simulations

The simulations given as examples here were run using the oxDNA code (June 2019 version). Structures were originally obtained in either Tiamat^12^ or CaDNAno^11^ format and then exported to oxDNA format using the TacoxDNA webserver^42^. Structures were relaxed in two steps. First, a brief Monte-Carlo (MC) simulation was performed using the DNA_relax or RNA_relax force fields to remedy overlapping particles. After this initial relaxation, mutual traps based on the intended design were applied to enforce relaxation to the intended design. For designs exported from Tiamat, mutual trap files are produced by TacoxDNA; for other structures, these files were produced using the force file generation script described in the “other utilities” section after the MC relax. A further relaxation was performed using the max_backbone_force option in a molecular dynamics (MD) simulation with the DNA2 or RNA2 force field. This bypasses checks of backbone bond length and allows for faster relaxation due to the CUDA implementation of the MD method in oxDNA^49^. This simulation was run until the energy stabilized between −57.98 and −62.13 pN nm (−1.4 and −1.5 oxDNA energy units respectively). At which point, the external forces and backbone force limitations were released and a production simulation run was performed using the same force field for 10^9^ steps with a stepsize of 15.15 fs (0.005 oxDNA time units). This corresponds to a total run time in the microsecond range; however, previous work with the oxDNA model^37,50^ suggests, in part, due to the increased diffusion coefficient, this direct conversion is an underestimate of the corresponding experimental time. However, as is the problem with all coarse-grained models, it is impossible to establish a direct correspondence between the simulation and experimental time because different processes in a coarse-grained simulation can scale to the experiment with different ratios. The production simulations were performed at 20°C using an Andersen-like thermostat^51^, and configurations were saved for analysis every 5 × 10^5^ steps, resulting in 2000 separate configurations used in each analysis. The RNA tile, used as an example of the clustering algorithm, was exported from a Tiamat design as described above. However, the production run was a parallel-tempered simulation run in virtual-move Monte Carlo (VMMC)^52^ with umbrella sampling. Simulations with eight parallel replicas were performed with the temperatures set from 25 to 60 °C at 5 degree increments. Replicate exchange was attempted every 1000 steps. To facilitate calculation of the energy barrier between the formed crossover and the broken crossover, a weight file was created to bias the simulation towards transition states between the two states.

## III. RESULTS

### A. OxView - Web browser visualization, analysis and editing of nanostructures

We introduce oxView, a JavaScript app built on the Three.js visualization library to provide fast, user-friendly, and flexible visualization capabilities with low technical overhead (Fig. 2). OxView uses hardware instancing to offload most calculation of object geometry to the computer’s GPU, allowing it to smoothly visualize structures containing millions of nucleotides (Fig. 2a). Standard Three.js scenes encounter a bottleneck in the rate of CPU draw calls with only a few thousand objects on the screen. By using instanced materials and custom properties written into the WebGL shaders, oxView bundles many objects with similar geometries into a single draw call that calculates edges and vertices in the compiled shader code.

**FIG. 2.**
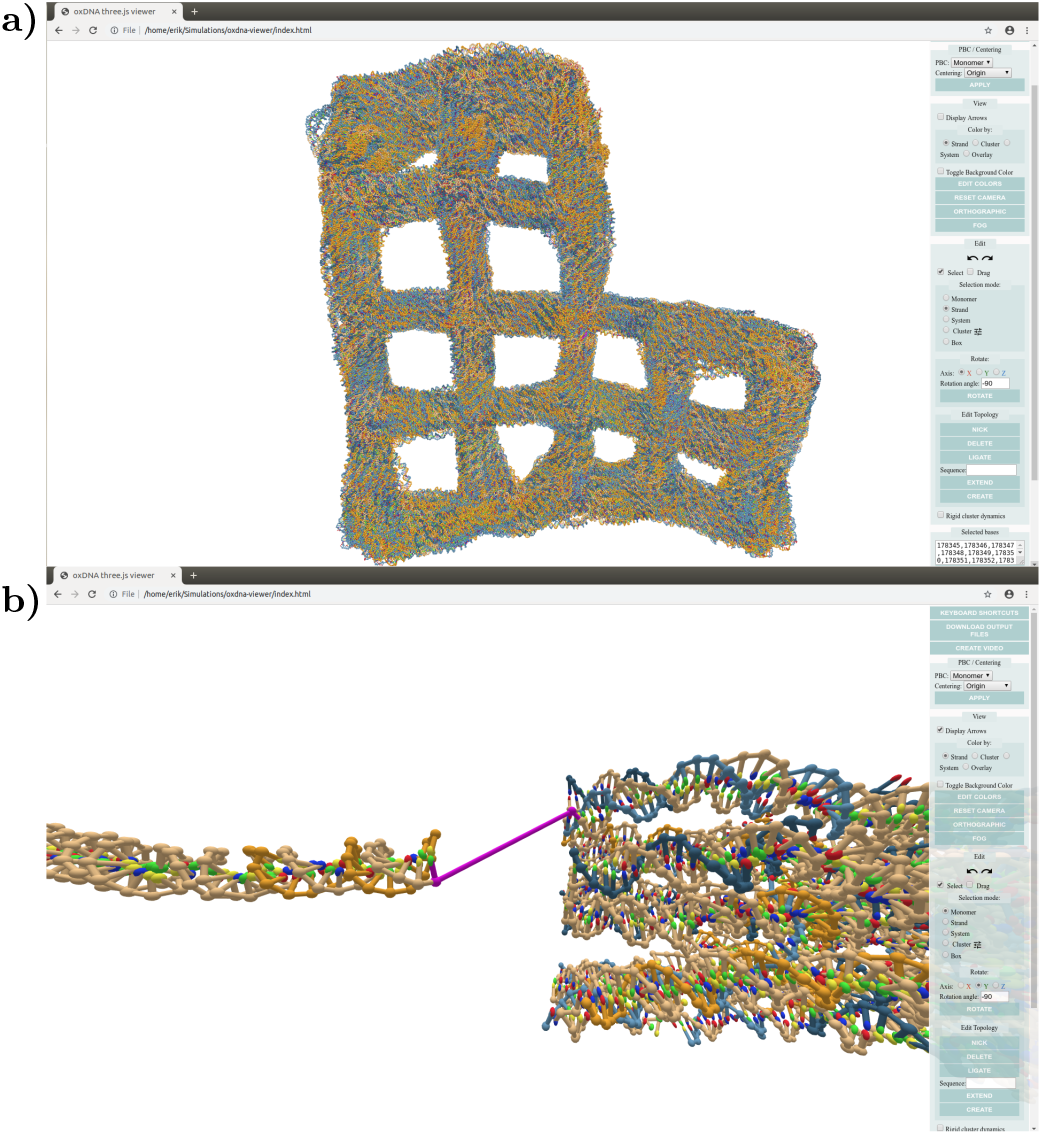
Screenshots from usage of oxView. **a)** 100 configurations from an oxDNA simulation of design 24 from^18^ merged into a single file and loaded into oxView; illustrating the ability to smoothly visualize over 10^6^ nucleotides. The origami design has 11 382 nucleotides, resulting in a combined file containing 1138 200 nucleotides, which renders as 5691000 individual objects in the scene. **b)** Using oxView to assemble a simulation of the tethered multiflourophore (TMF) structure used in^54^. Each of the subunits is a separate CaDNAno file converted into oxDNA format using^42^. The two subunits and the algorithmically generated tether had to be ligated prior to simulation.

Loading a simulation is as simple as dragging and dropping a trajectory/topology file pair onto a browser window with the app running. OxView will run in any modern web browser. Though, we note that Google Chrome tends to have the best performance for very large structures due to a better WebGL implementation. Simulation trajectory files can be stepped through using onscreen buttons or the keyboard, and the trajectory movie can also be downloaded as a video file. Available formats are .webm, .gif, and .jpg/.png image archives.

In addition to visualization, oxView also has basic editing capabilities (Fig. 2b). Particles can be selected individually, or whole strands and systems can be selected as a whole. Box selection, range selection (shift+click) and cluster selection are also available. Clustering can be done automatically using a Density-Based Spatial Clustering of Applications with Noise (DBSCAN) algorithm^53^ or can be assigned manually from other selection methods. Briefly, DBSCAN compares distances between points and classifies groups of points meeting a specified minimum size and within a specified mutual minimum distance as members of the same cluster. It also characterizes points as central or peripheral, with central points having at least the minimum number of neighbors in the cluster and peripheral points being within the cutoff distance of one or more, but fewer than the specified number of central points. When manually selecting nucleotides, holding the control key while making a selection will combine the new and previous selections. Selected particles can be translated and rotated, and the topology can be edited via strand extension and creation, nicking, deletion, and ligation. Edits can be undone and redone using the standard ctrl-z/ctrl-shift-z keyboard shortcuts. Strand extensions will attempt to approximate either an A-form or B-form helix depending on the parent nucleotide’s identity: RNA or DNA. The final edited version can be downloaded as an oxDNA file pair for further simulation or as a CSV sequence list for experimental validation.

We envision this tool being used to prototype DNA/RNA nanostructural designs in an iterative process before realization in the lab. The structure can be simulated for a short time, analyzed for defects, and then iteratitively modified in the viewer and returned to simulation to verify success. This tool is also useful as a neutral ground between structures designed in other editing tools, allowing researchers to merge together structures from many sources to realize a complex vision.

oxView also allows the creation of mutual trap external force files for oxDNA/RNA. These files define artificial pairwise spring potentials between nucleotides that can be loaded in an oxDNA simulation and be very helpful when simulating the relaxation of a complex structure, assembled from multiple components, or when relaxing a structure imported from the CaDNAno format.

#### 1. Implementation details

The underlying architecture of oxView has two parallel data streams. The first mirrors the physical arrangement of nucleic acid monomers into strands, with each configuration/topology pair representing a system. This data structure contains the topological information relating to particle identities, connectivity, and relation to the system. Monomers, strands and systems all inherit from the Three.js Group object and are related through an inheritance hierarchy, which allows interaction with structural units as a group. Additionally, each system contains a set of data arrays that define the positions, orientations, sizes, and colors of every particle. These arrays are passed into a custom implementation of the WebGL Lambert shader, where they are compiled on the GPU and drawn as a single object. This scheme allows loading of over 1 million nucleotides into a single scene (Fig. 2a and Supplementary video 1).

Selection is handled through a GPU-picker, which avoids the need for computationally-expensive raycaster intersection calculation. Briefly, each nucleotide has a mesh with a color corresponding to its global ID at the same position as its backbone site which is rendered in an invisible scene. The color of this mesh can quickly be determined via the x-y coordinates of the mouse on the screen. When the color is converted from the hexadecimal color to the corresponding decimal value, it returns the ID of the nucleotide under the mouse pointer. As the arrays passed to the shader are of constant-size, new nucleotides added to the scene after initialization, are placed in a temporary system object with its own instancing arrays.

Editing a simulation topology via nicking, ligation, addition, and deletion can leave gaps in the strand and nucleotide numbering. Because of how oxDNA allocates memory, such gaps will cause exported simulations to crash. Therefore, prior to exporting files from scenes with edited topology, the “pre-organize strands” option must be selected in the ‘‘Download Output Files” dialog (illustrated in Supplementary video 2). This option will re-organize strands from longest to shortest, ensuring that all numbering is contiguous, and oxDNA will be able to read the edited file. This also reinitializes the data arrays, merging the scene into a single system.

#### 2. Data overlays in oxView

Many of the simulation analysis scripts introduced in this work output overlay files that can be viewed in oxView. This allows interactive visualization of different properties (such as flexibility, discussed in Fig. 4) of respective parts of the structure obtained from simulations. These are JSON-format files that define the name of the overlay and the data. There are three types of overlays recognized by oxView. The most frequently used is the color overlay. These files contain one value per particle. When dragged and dropped into oxView, along with the corresponding configuration/topology pair, the color overlay file will create a superimposed colormap on the structure based on the value associated with each particle. All 256-value colormaps from Matplotlib^44^ are available in addition to the default Three.js colormaps. The displayed colormap can be altered via a simple API implemented in the browser console. In addition to per-nucleotide coloring, oxView can also read two JSON formats corresponding to arrows drawn on the scene. The first is a three-component vector for each nucleotide, which is produced by the principal component analysis script and draws a vector, emanating from each particle, using the magnitude and orientation defined in the overlay file. The second format, which can contain any number of vectors, takes pairs of three-component vectors and draws arrows of the corresponding position and orientation on the scene.

#### 3. Relaxing structures using rigid body dynamics

There has been a recent push to develop software that converts structures designed in the various design tools to simulation formats^42^. Due to the lattice-based drawing platform with parallel helices used by CaDNAno, exported structures can be very difficult to relax to a physically reasonable state in oxDNA. Initial configurations imported from CaDNAno (shown in Fig. 3a) will generally be planar with highly stretched bonds between individual structural units. Thus, without 3D information on how to reorient the helices, neither MC nor MD simulations are able to find the relaxed arrangement. This can also lead to topological impossibilities, where structures are knotted in a nonphysical manner. Additionally, starting simulations from a state with very stretched bonds can result in numerical instabilities that crash the simulation. For origami structures consisting of multiple origami blocks, connected by initially stretched backbone bonds, rigid-body manipulation has previously been used to arrange the converted oxDNA structure into a more realistic initial configuration^55^. The translation and rotation tools in oxView allow users to select and rearrange blocks of nucleotides as rigid bodies. Furthermore, oxView also includes a rigid-body dynamics (RBD)^56^ mode, that automatically transforms groups of nucleotides based on a simple force field. It is also possible to drag and rotate groups during RBD, allowing the user to nudge the design into the desired topology. Groups can either be created manually via the selection interface or through the implemented DBSCAN algorithm^53^ that automatically identifies and categorizes spatially separated groups of particles. The latter option works particularly well with designs developed in CaDNAno.

**FIG. 3.**
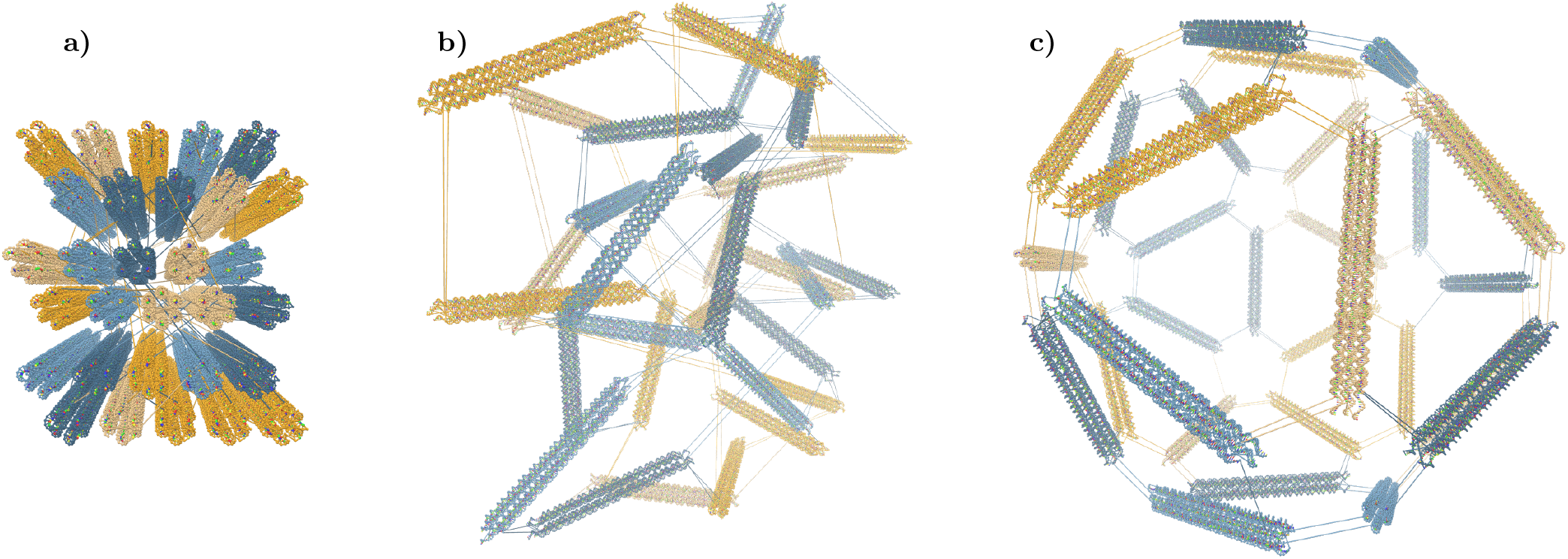
Rigid-body dynamics of clusters. Snapshots from the automatic rigid-body relaxation of an icosahedron, starting with the configuration converted from caDNAno **a)**, through the intermediate **b)** where the dynamics are applied, and **c)** the final resulting relaxed state.

Each group is represented as a rigid body with a position and an orientation. The groups are held together with spring forces at each shared backbone bond, with a magnitude of

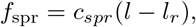

where *c_spr_* is a spring constant, *l* is the current bond length and *l_r_* is the constant relaxed bond length. To avoid overlaps, a simple linear repulsive force, of magnitude

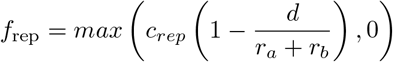

is added between the center of each group, where *c_rep_* is a repulsion constant, *d* is the distance between the two centers of mass, and *r_a_* + *r_b_* is the sum of the group radii (the greatest distance they can be while still overlapping). An example of the dynamics in action can be seen in Fig. 3 and Supplementary video 3, where each side of a DNA icosahedron^57^ is automatically arranged into the intended shape.

### B. General-purpose analysis tools

Popular molecular simulation tools programs, such as GROMACS^58^, not only perform molecular simulations, but also include analysis tools for common use-cases. The access to reliable and maintained tools, as part of the distribution, allows for standardization between many researchers using the core tool, as well as simplifying the learning curve for new researchers working with the tool. At this time, although there are over a hundred publications using oxDNA/RNA, no standardized set of tools for structural analysis has emerged. We present here a set of tools covering many common structure analyses: mean structure, root mean squared fluctuations (RMSF), hydrogen bond occupancy, interaction energy, interduplex angles, contact mapping, the distance between nucleotides, and principal component analysis of structure motion. These are primarily written in Python, with some portions embedded in the oxDNA C++ code for enhanced speed. Moreover, we provide additional utilities including a parallelization scheme for analyses, trajectory alignment, and unsupervised clustering based on data outputs.

#### 1. Mean structure determination and RMSFs

This package includes two methods for determining the mean structure. One utilizes the Biopython^45^ singular value decomposition (SVD)-based structure superimposer. This is a popular method^59^ that finds a translation and rotation to superimpose two distinct conformations on top of each other to minimize the the root mean square distance between their components. A random configuration in the trajectory is selected as the reference structure. In the example structures displayed here, this choice was found to have little impact on the final outcome. Each configuration is then superimposed onto the reference, and the average position of each nucleotide is calculated by taking the mean of each particle’s coordinates in the aligned reference frame. To find the per-particle RMSF, a second script uses the mean structure produced by the first script as the reference configuration for alignment. The squares of the distances between the alignment and the mean structure for each nucleotide are then summed and divided by the total number of configurations. The square root is then taken to find the RMSF per particle in nanometers. The final output from this script is a .json format color overlay that can be loaded into oxView.

As noted in^23^, averaging methods that use full structure alignment work very well for rigid structures; However, there are some caveats. Large planar structures frequently appear to have the smallest RMSF in a ring midway between the center and the edge (Fig. 4b). This does not correspond to lower flexibility, but instead reveals an artifact of the single-value decomposition. If a structure can bend in two possible directions, the stationary point in the oscillation will appear to have very low flexibility. Highly flexible regions tend to collapse towards a center line, which is particularly problematic for rigid structures connected by a flexible linker, exemplified by the interrupted duplex shown in Fig. 5a. When the average structure is computed for this design, the entire structure collapses into a linear blob that does not have any resemblance to any of the individual configurations. This is because the average position for these flexible particles is drawn towards the center. For such structures, another mean structure calculation based on interparticle distance is employed.

**FIG. 4.**
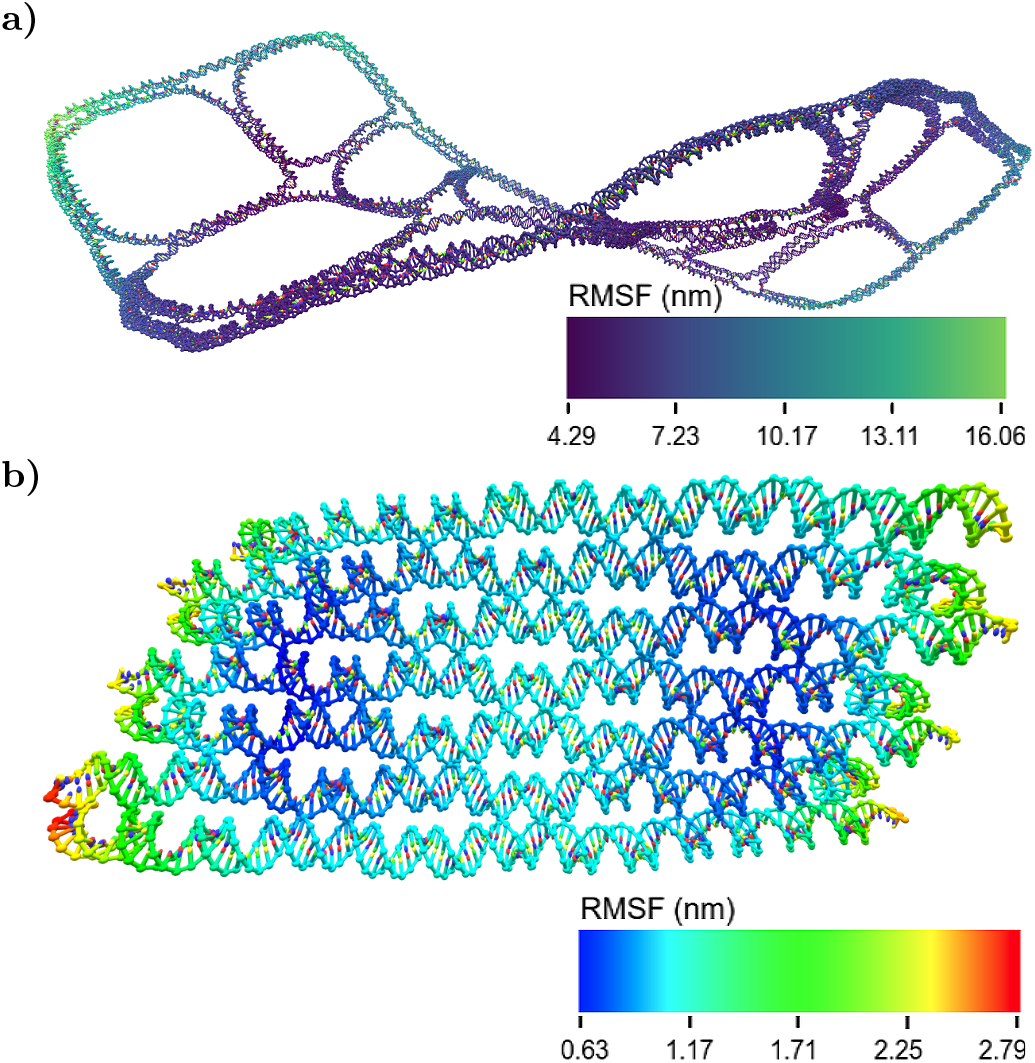
Mean structures and RMSF. **a)** The mean and deviations scripts were used to compute the mean structure and RMSFs of design 19 from^18^. In the initial report of these designs, they were characterized by AFM, showing complete, flat structures. In the simulations here, the structures were stable; however, the mean structure shows a significant right-handed global twist. **b)** To demonstrate the patterns that appear in RMSF calculations, this is the mean structure of a single-stranded RNA origami^65^ with the RMSF shown using a colormap with high spectral contrast. The center of the origami appears to have an RMSF twice as high as the surrounding regions. This is simply an artifact of the alignment and not an accurate characterization of particle motion.

The second option for mean structure determination uses a common machine learning technique, multidimensional scaling (MDS)^60^, to reconstruct a mean structure from local contact maps. MDS is one of a class of algorithms known collectively as manifold learning, which are traditionally used to perform dimensionality reduction in high-dimensional datasets. MDS takes a set of pairwise distances between points in an arbitrary number of dimensions, as an input. The algorithm then uses eigenvalue decomposition to find distances *d_ij_* in the embedded space that minimize

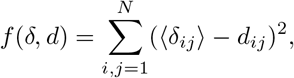

where N is the number of data points, 〈*δ_i,j_*〉 is the mean distance between centers of mass of nucleotides *i* and *j* (averaged over the whole simulated trajectory) and *d_i,j_* is their embedded distance^46^. In the implementation presented here, pairs of nucleotides, where average distance 〈*δ_i,j_*〉 is longer than the cutoff of *r*_cut_ = 2.07*nm* (approximately the interhelix gap in an origami), are not considered in the embedding. The MDS-based mean structure calculation uses the MDS algorithm^61^, implemented in the Python machine learning toolkit, SciKit-Learn^46^, to reconstruct these local distances into a three-dimensional embedded representation. This method loses orientation data, and thus, nucleotides are simply visualized as spheres at their centers of mass (Fig. 5). Once a mean structure (in the embedded space) is calculated, the script then calculates the mean deviation in distance between each particle and its nearest neighbors and outputs an oxView color overlay file to quantify the flexibility.

**FIG. 5.**
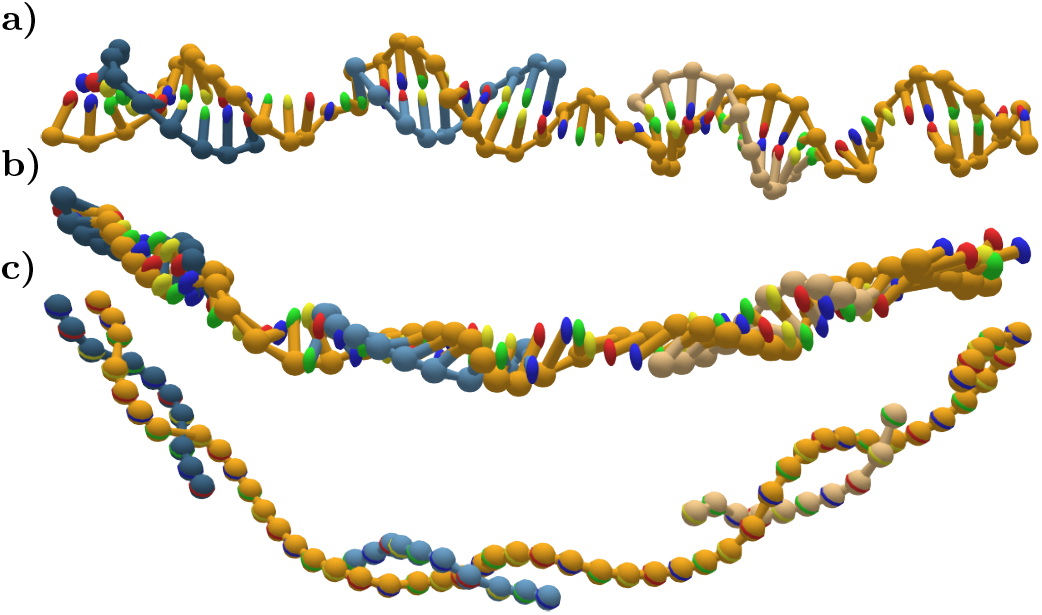
Improving mean structures of flexible designs. **a)** The initial configuration of a 50-nucleotide duplex interrupted with 5 nucleotide gaps, created using the editing tools in oxView. Each individual configuration encountered during simulation displayed helical geometry. **b)** The mean structure computed using SVD of the whole simulation. Because of the high backbone and rotational flexibility of this structure, it collapses into a linear shape that has little correspondence to the double helix geometry that is maintained throughout the simulation, **c)** The mean structure computed using MDS. In this case, since only local contacts are used to construct the mean structure, the helical geometry is maintained. MDS comes at the cost of losing nucleotide orientation information, however. Thus, the visualization only shows the center of mass for each nucleotide.

We used the SVD-based mean structure script to study flexibility and curvature in large wireframe origami structures^18^. In the original research, these structures were visualized using atomic force microscopy (AFM), which tends to overestimate the flatness of structures due to electrostatic interactions between the mica surface and the DNA origami^4^. Though the wireframes appear flat in the published AFM results, our simulations suggest that in solution they would be more crumpled or have some degree of global helical twist. Particularly striking is the helical shape of the mean structure of design number 19 from^18^ (shown in Fig. 4a and Supplementary video 4). OxDNA was parameterized to correctly reproduce the global twist of large 3D DNA structures^48,62^, suggesting that this twist is likely significant while in solution. We note, however, that the global twist of 2D DNA nanostructures in the bulk remains a topic of active research^63^, and more experimental data is needed to establish a better comparison of oxDNA parametrization with experimentally determined structures. Mean structures are also the best method to compare simulation results to cryo-EM maps. Both produce an averaged structure over thousands of individual snapshots. Thus, converting mean structures to PDB format using existing conversion tools^42^ for use with cryo map fitting software, such as can be found in Chimera^64^, is a method to correlate simulations and experimental data.

Because of the limitations of SVD-based mean structure calculation, the MDS approach was also used to determine the mean structure and deviations. Unfortunately, because average distance data is noisy and does not precisely map to a single configuration, this method does not work for structures larger than a few thousand particles. In all tests of the algorithm at origami scales, every particle was placed at the origin, a trivial solution that is a known issue of manifold learning methods. However, at smaller scales, this method provides a reasonable mean structure, that respects the geometry of the double helix, and a measure of deviation that reveals areas of flexibility without global artifacts due to fitting (Fig. 6).

**FIG. 6.**
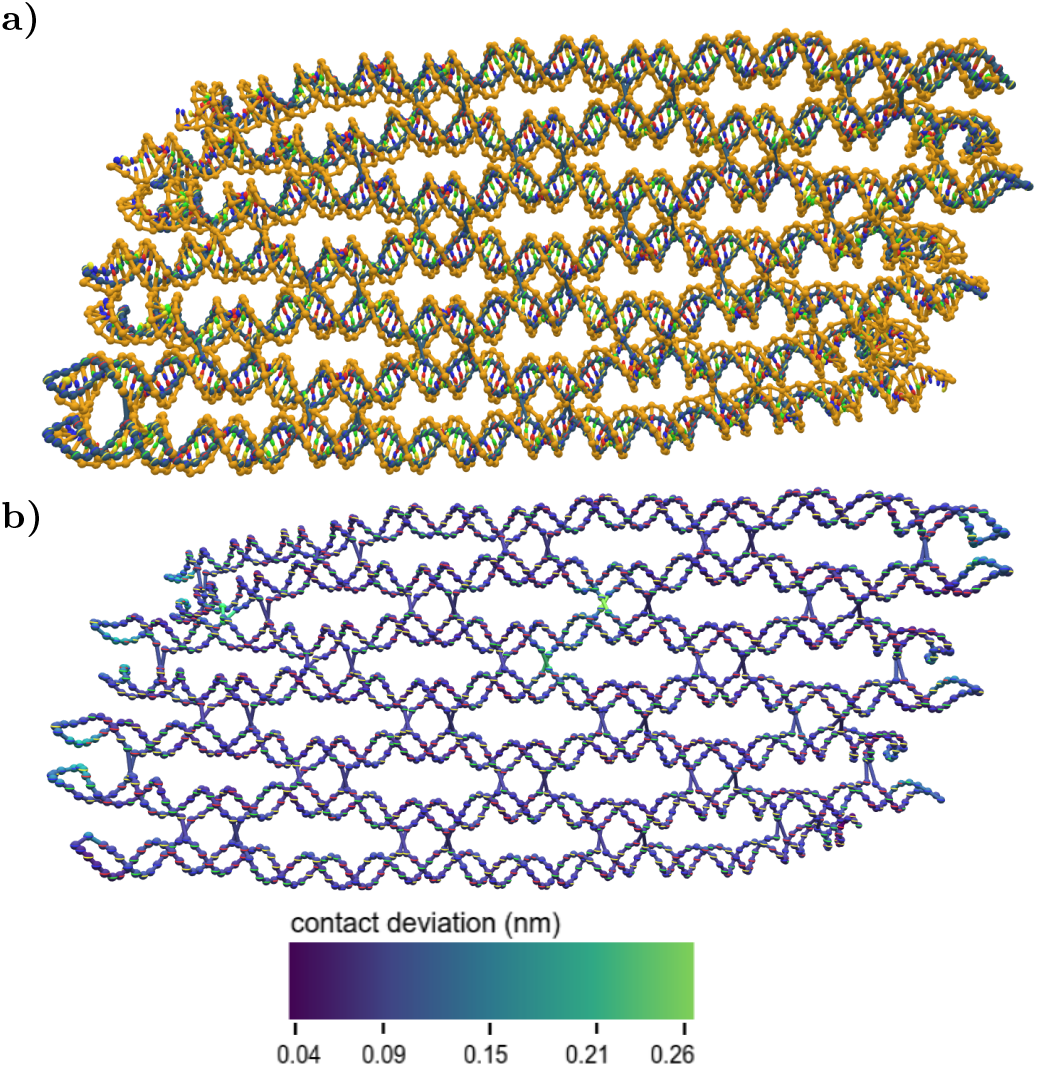
Mean structures computed via multidimensional scaling. **a)** The mean structure of a single-stranded RNA origami from^65^ computed both by SVD (yellow) and MDS (blue). Because MDS does not preserve orientation data, the nucleotides are visualized simply as spheres at their center of mass, rather than having distinct base/backbone sites. **b)** The deviation in local contacts from the mean structure calculated in **a**. This measure shows most of the structure to be homogeneously stable, with higher flexibility at helix ends and at junctions capable of sliding.

#### 2. Geometric parameters: interduplex angles and distances

The simplest structural unit of nanotechnology structures is the duplex - antiparallel strands of sequentially bonded nucleotides. We have implemented a script that automatically determines the duplexes present in each configuration within a trajectory and fits a vector through the axis of the duplex. This is trivial for DNA, where the center points of each base pair lie roughly co-linear and the axis can be defined by a linear regression through the points in the center of the duplex. For RNA, the A-form helix is slightly more difficult to characterize. The duplex is defined by the normal vector to an average plane fit through the displacements along the backbones as described in^31,66^. This script creates a text file that contains information about all duplexes found at each step. This can be visualized using a separate script, which uses the ID of nucleotides at the edge of the duplex, found using oxView’s selection feature. This method can compare angles either within or between structures.

Determining the angle between two duplexes can be useful in assessing design outcomes as well as quantifying twist within nanostructures. The output from the angle script is a list of all duplexes found in each configuration of the trajectory. This output can then be fed into the partnered visualization script along with the starting nucleotide IDs of the duplex. The output will be the median, mean and standard deviation of the angle between the two duplexes, as well as the fraction of analyzed configurations in which that pair of duplexes are both present. This number is an indication of both how stable the structure is and whether or not the chosen duplex is representative of the entire trajectory. The script will also provide a histogram and/or trajectory of the angle over the course of the simulation. Here, we show an example of the angle script again using the wireframe origami designs from^18^. Each origami has a designed junction angle corresponding to the number of arms joined at each junction (Fig. 7). Deviation from this designed angle is a measure of strain and how non-planar the structure is in simulation. This can be particularly revealing in combination with the mean structure, showing that an on-average flat structure has a significant degree of flexibility over the course of the simulation.

**FIG. 7.**
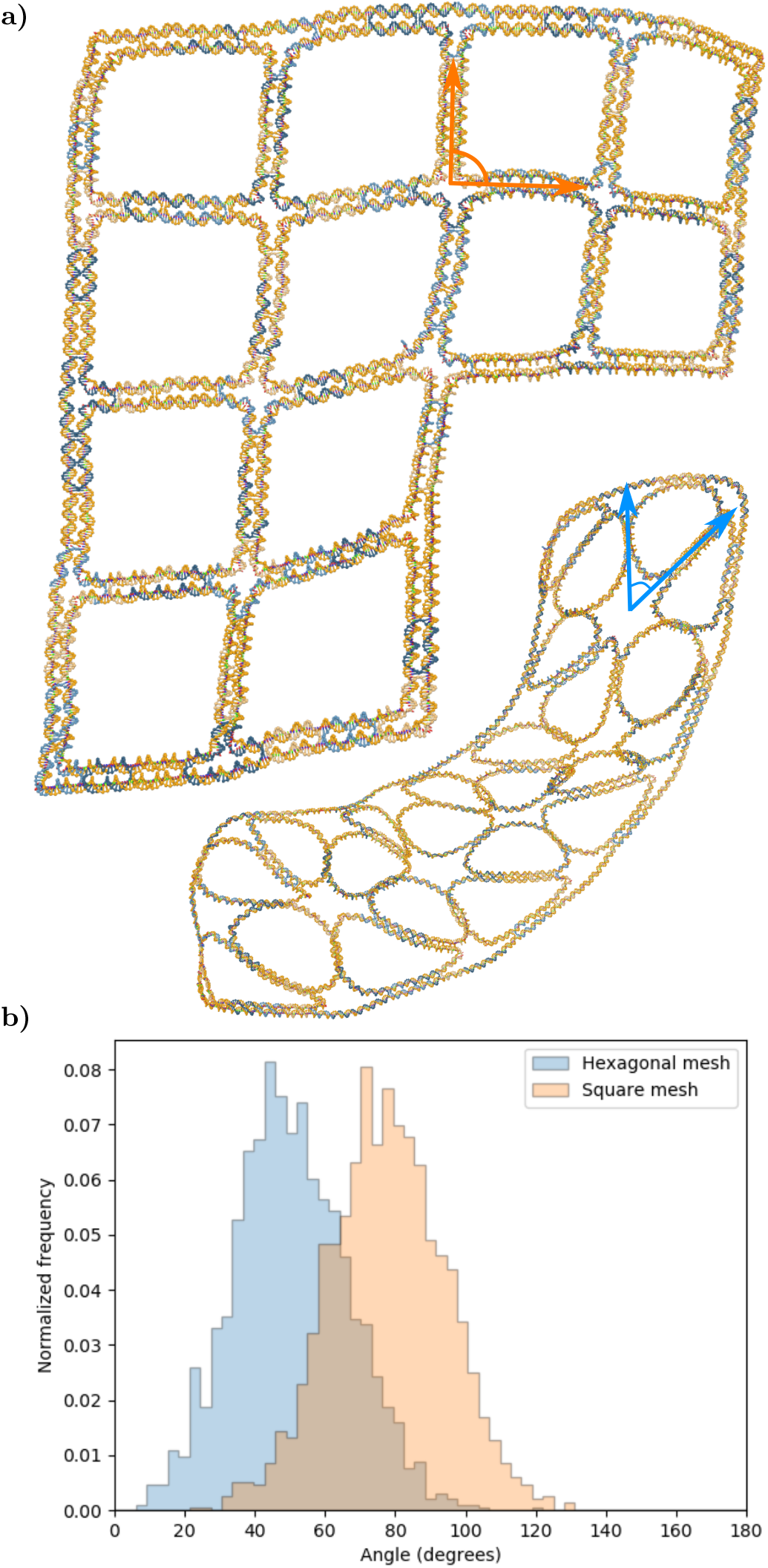
Comparing angles in wireframe lattices. **a)** The mean structures of design 23 (top) and design 20 (bottom) from^18^. The structures are designed to have a square and hexagonal lattice pattern, respectively. **b)** The distribution of angles between two arms of a junction showing variation around the designed junction angle. For the hexagonal lattice, the observed angle is lower than the designed angle of 60° because the structure has significant out-of-plane curvature in the simulation.

The Tethered Multi-fluorophore (TMF) structure from^54^ was used as a demonstration of the distance script. This structure is used to measure binding kinetics through the large change in radius of gyration induced by binding and unbinding of compatible sequences near the ends of the double-stranded tether. Fig. 8 shows end-to-end distance of the tether in both the bound and unbound states. Knowing the end-to-end distance of this structure can be used in predicting the radius of gyration for various states of the structure, which is useful in corroborating experimental results.

**FIG. 8.**
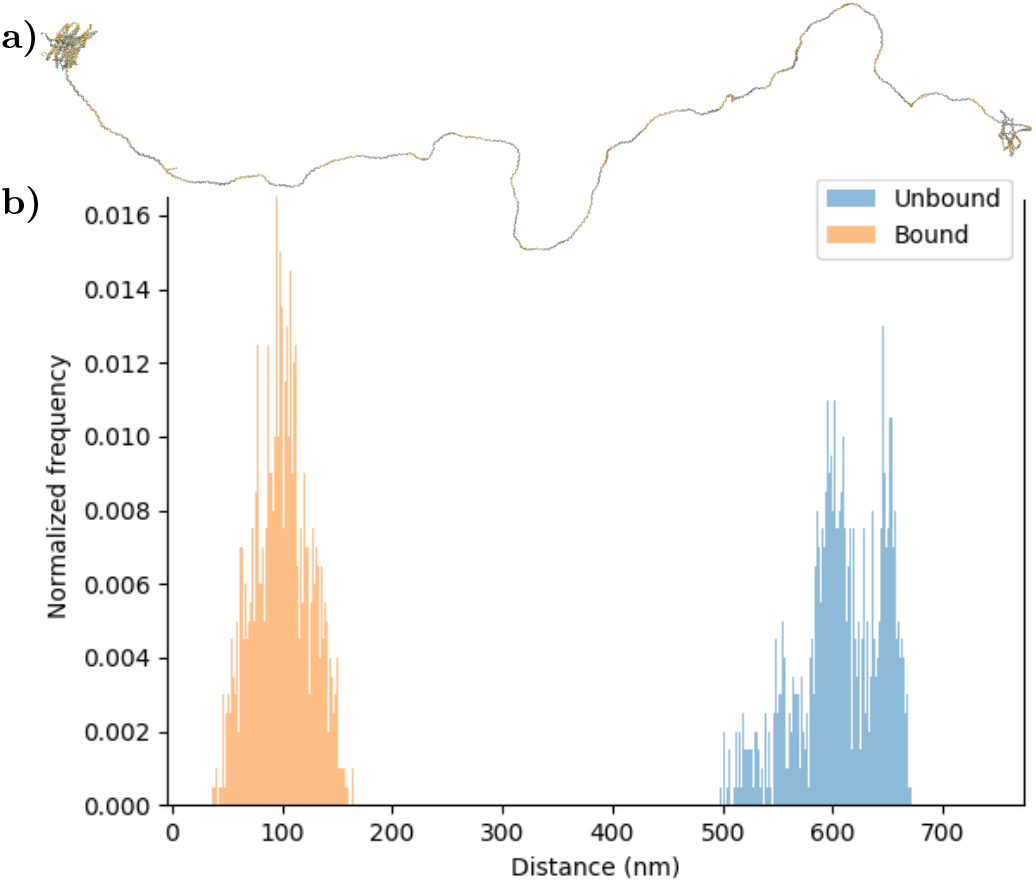
Distance between origami units of TMF. **a)** The final configuration from a simulation of the TMF structure used in DNA kinetics experiments^54^. Separate simulations were performed with the sticky ends in both the bound and free configurations. **b)** The distribution of distances between the origami units at opposite ends of the tether.

#### 3. Base pair occupancy

The hydrogen bonds defining Watson-Crick base-pairing are the single most important parameter defining DNA/RNA nanotechnology geometries. Since structures are designed towards a theoretical global free-energy minimum that maximizes hydrogen bonds, deviations from the designed structures point to regions of significant topological strain or that have found a kinetically trapped structure distinct from the intended design. OxDNA/RNA defines hydrogen bonds between base-paired nucleotides as a base-pairing potential between two base particle beads less than −0.1 k_b_T, about 10% of the magnitude of the equilibrium value of the base pairing potential of a base pair in a duplex. The script compares the hydrogen bonds in a simulation with a provided list of pairs present in the intended design. The fraction of the configurations in which the intended bonds are formed are reported as an oxView overlay file, with color coding intensity corresponding to the fraction of the time where the bonds are formed. Bonding is considered 0 for nucleotides without designed complements.

Since the structures exported from design tools represent an idealized form, deviations from the original vision imply unmet design constraints. In Fig. 9, we use this script to explore a poorly-formed RNA tile structure. We first simulated the original tile design, as shown in Fig. 9a. The hydrogen bond occupancy data revealed intense stress in a single duplex, with individual bonds ranging from 0-60% occupancy. This introduced considerable flexibility to the structure, disrupting the intended planar design. When the duplex was redesigned to extend it by one base pair, it no longer suffered from the same disruption, and the intended design was observed in the simulation (Fig. 9b).

**FIG. 9.**
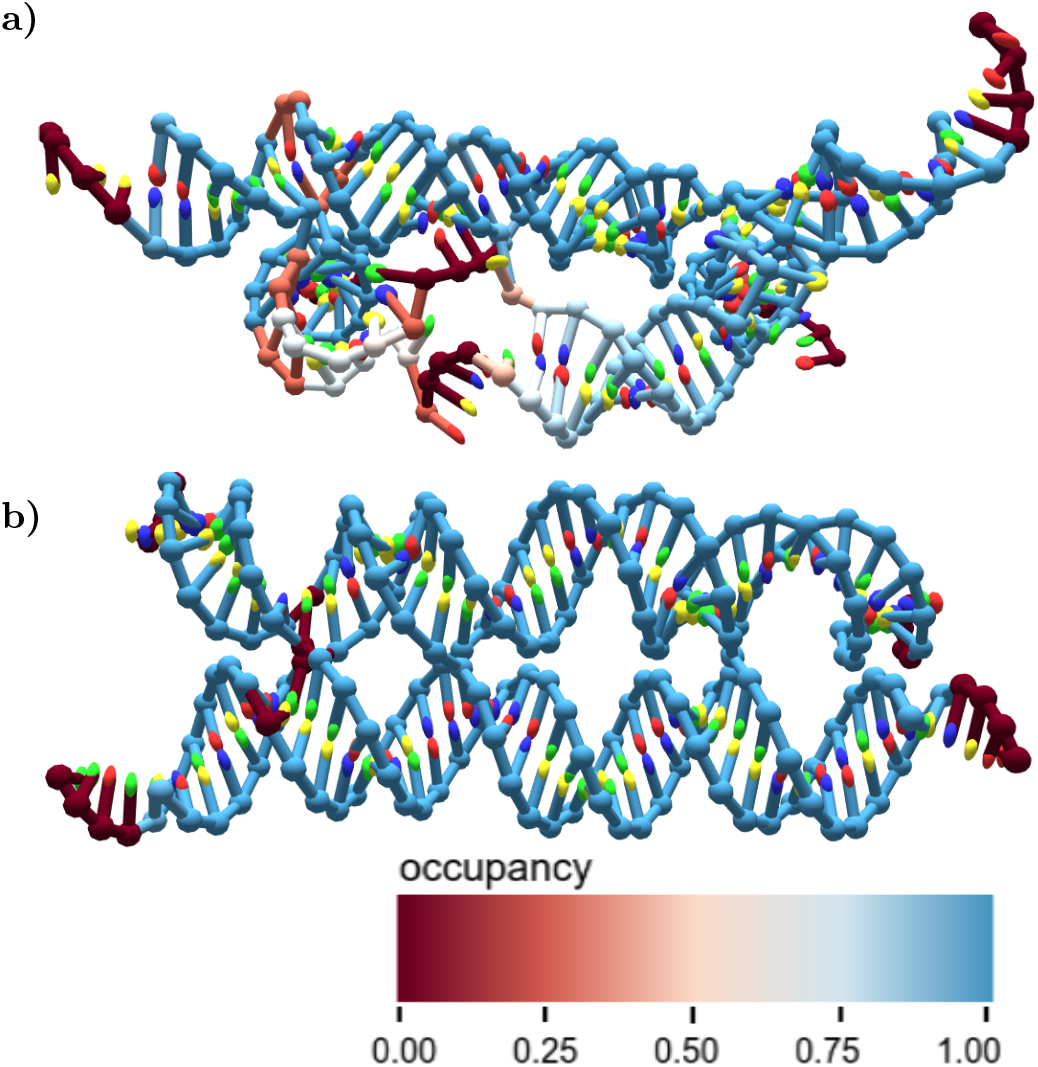
Bond occupancy of an RNA tile. **a)** The hydrogen bond occupancy during an oxRNA simulation, overlaid on a structure of an RNA tile. The structure was known to form poorly in the lab, and the simulation revealed significant strain on one duplex. The structure used here is the centroid of a trajectory based on the global fitting parameters discussed later. This was used as a visualization instead of the mean structure, as the unpaired duplex made the structure so flexible that the mean structure collapsed. **b)** The broken duplex from the structure in **a** was extended by one base pair, and the simulation was re-run. Shown here are the hydrogen bond occupancies overlaid on the mean structure. In simulation, this significantly improved rigidity.

#### 4. Principal Component Analysis of Nanostructure Motion Modes

Principal component analysis (PCA) is a common method for analyzing molecular simulation data that extracts the largest sources of deviation from the dataset^67^. First, using SVD, each configuration is aligned to a mean configuration (produced by either SVD or MDS) to remove rotations and translations from the data. Each nucleotide’s deviation from its reference position in x- y- and z-coordinates is stored as its difference matrix. A covariance matrix is then constructed from the difference matrices, and the eigenvalues and eigenvectors are found through eigenvalue decomposition. These are then sorted in descending order with the highest eigenvalues representing the largest sources of variation in the structure. The eigenvectors generated by PCA represent an orthogonal basis for the reconstruction of every structure visited during the trajectory, and these reconstructions can then be used for clustering of distinct sampled conformations. Finally, the PCA script outputs a .json file for the oxView tool, which displays arrows on the structure corresponding to the sum of a user-defined number of components weighted by their respective eigenvalues.

To demonstrate the principal component analysis of DNA/RNA structures developed in this work, we ran it on a simulation of a Holliday junction (Fig. 10). As one would expect for this structure, PCA reveals strong collective motion for the junction arms. The motion grows stronger at the ends of duplexes, while the crossover point shows little motion.

**FIG. 10.**
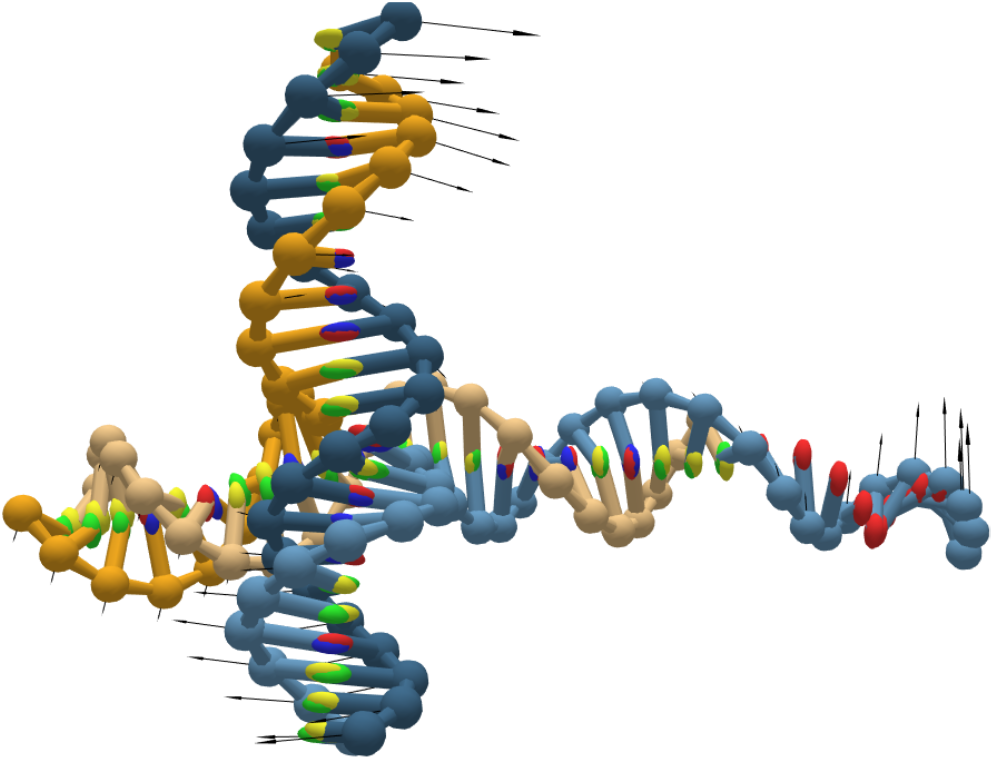
Principal Component Analysis of a Holliday junction. Principal component analysis of a Holliday junction visualized on oxView. Shown here is the top mode, which corresponds to a scissoring motion in the junction, with the arm ends having significantly higher average displacement than the crossover point.

#### 5. Unsupervised clustering of configurations encountered in simulation

The trajectories produced in an oxDNA/RNA simulation can be tens of gigabytes in size and explore an expansive amount of the configuration space available to the structure. In cases where multiple metastable states are visited during the trajectory, aggregate structural data, such as mean structures or base pair occupancy, might not be representative of the ensemble. This is due to the presence of these distinct metastable states. Here, we once again use the DBSCAN clustering algorithm^53^, as implemented in SciKit Learn^46^, to automatically extract geometrically distinct structures from large trajectories and save them as new trajectory files that can be analyzed independently. The clustering algorithm can take any matrix of positions as an order parameter, whether that be principal component coefficients of each configuration, or simply the distance between two particles. The DBSCAN algorithm is particularly good at clustering molecular simulation data where metastable states tend to form distinct clusters separated by a large energy barrier, such that observing transition states is relatively rare and multiple distinct densities are observed.

To demonstrate the utility of clustering using structural order parameters, we analyzed a simulation of an RNA tile structure (Fig. 11), that is known to form two distinct structural isomers in experiment (unpublished results). In the simulation, two states were encountered, the correctly-folded structure, with three crossovers, and an unfolded structure, in which the paranemic cohesion^68^ between two of the crossovers is lost, leaving essentially a Holliday junction (Fig. 11). There are many potential order parameters that can be used to separate out these two structures. In this case, we chose to work with the most aggregate data: each configuration’s position in principal component space.

**FIG. 11.**
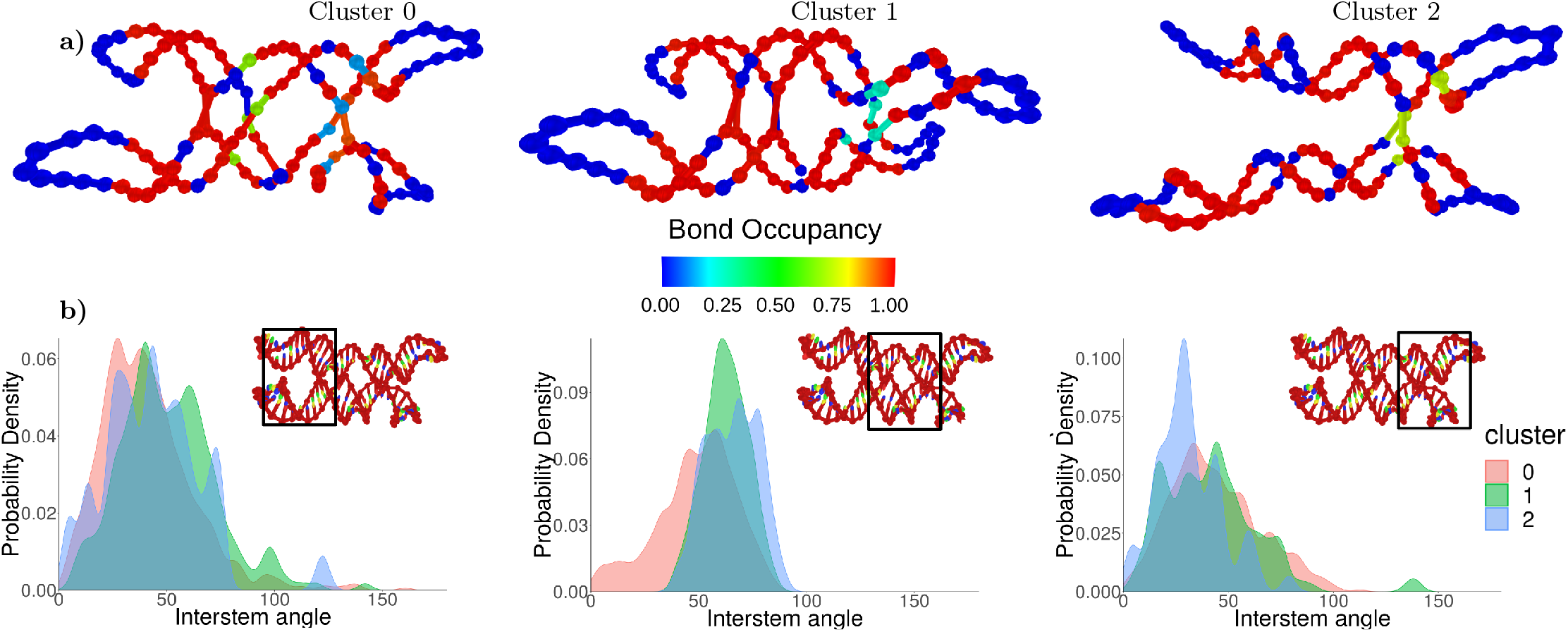
Unsupervised clustering to isolate isomers of an RNA tile. **a)** The three clusters found in a simulation of a single-stranded RNA tile. The mean structure of each cluster was determined using MDS, and the hydrogen bond occupancy compared with the original design was used as an overlay. **b)** Histograms of the angles found in each cluster showing the distinct structures found in each cluster. The black frame on the tile snapshot indicates pairs of double-stranded RNA regions that were used to calculate the interstem angle.

The components produced by PCA represent a linearly independent basis for describing structures relative to the provided mean structure. This also means that every configuration used to compute the components can be mapped to a unique point in 3N-6 dimensional space. When applying DBSCAN to the positions of configurations in this space, the distinct conformational isomers can be separated without further processing. In addition to the two expected configurations, this method also separated out another cluster (cluster 1 in Fig. 11) of structures where the paranemic cohesion was correctly formed, but stacking was interrupted at the nick point, resulting in a non-planar kinked structure. The clustering script automatically produces separate trajectories for each of the identified clusters; these were further analyzed using the angle script, identifying the distinct interduplex angles between each duplex in the structures (Fig. 11b).

#### 6. Other Utilities

In addition to the specific structural measures discussed here, this package also contains additional utility functions for processing and displaying data. The first are two scripts that utilize the SVD superimposer from Biopython^45^ for improving visualizations. The superimposing script takes multiple configuration files that share the same topology and returns them with their translations and rotations removed relative to the first configuration provided. We find this very helpful for comparing mean structures of similar designs or of the same design under different simulation conditions. There is also an alignment script, which takes a trajectory file and aligns all configurations to the first one in the file. This makes for a much smoother visualization experience when exploring trajectories in oxView or when making movies of a trajectory.

We have found the alignment scripts to be very useful for producing figures and movies (see Supplementary video 5 and Fig. 6a) and for making comparisons between designs. These scripts are limited, however, by the need to align discrete units. Therefore, the structures must have the same number of particles in mostly the same position. Thus, the scripts are best used for comparing simulation conditions, changing sequences, and changing crossover positions in designs.

There is also a utility that reports the energy contribution of every interaction in the model. This has options of a text output to check specific values, as well as an oxView overlay showing the average energy of all nucleotides over the course of a simulation. Checking the base pairing or stacking interactions of specific nucleotides can be very helpful in identifying properties or defects in a given design. Additionally, we have found the visualization option useful for identifying excluded volume clashes during relaxations of large structures, as these cause extremely high total energies, which visually pop in oxView.

There are two further scripts that work with base pairs. One takes the current arrangement of base pairs in the structure and generates either the designed pairs file used by the base pair analysis script, or an oxDNA mutual trap force file, which can be used to enforce a particular base pairing configuration during relaxation. This can be particularly helpful when relaxing multi-component structures edited in oxView, as the forces pulling stretched bonds back together can cause unwanted fraying of base pairs in otherwise stable structures. The second script converts oxDNA force files into a designed pair file. The Tiamat converter from^42^ can produce force files as part of the conversion process, and this script can convert those force files into the format needed for the duplex angle script.

Finally, we provide a parallelization scheme for analyzing oxDNA trajectories. The parallelization module breaks down a trajectory into a number of chunks equal to the number of CPUs you have available, and uses the Pathos Multiprocessing library^47^ to map trajectory chunks, CPUs, and functions. If the user has enough computational resources available, this facilitates analysis of even very large structures or long trajectories in a matter of minutes. The implementation of parallel functions is standardized across all scripts used here, and users are encouraged to follow the example given here in developing further analyses specific to their own designs.

Most of the analysis discussed fall into the class of tasks known as “embarrassingly parallel”, where there is no communication required between processes, and the final joining step is relatively easy. For all structure analysis algorithms described here, each configuration can be calculated independently of all the others. The only limitations to parallelization come from calculating split points in the trajectory and if a data trajectory is required, combining the outputs together in the proper order. As an example, we benchmarked parallelizing the computation of the mean structure of two structures: one with 423 nucleotides, and the other with 11385. In both cases, runtime decreased by more than a factor of 10 when run on 30 CPUs compared with a single CPU, with diminishing returns past that point.

## IV. DISCUSSION

We developed this collection of tools to remedy two gaps that we have perceived in the oxDNA software environment. First is the lack of an all-in-one visualizer that loads files within a reasonable timeframe, has a user-friendly UI, and performs edits on structures that could then be further simulated. All-atom simulations have such tools in the form of VMD, Chimera and PyMol. While tools exist to convert between all-atom and oxDNA formats, this is a cumbersome process that we felt could be remedied by the development of oxView. The use of hardware instancing allows oxView to load structures of unprecedented sizes and facilitates our work on million-nucleotide oxDNA simulations of multi-origami structures. Furthermore, because oxView is built using the open-source 3D library Three.js, opens the possibilities for features from other Three.js projects to be added to oxView. For example, virtual reality oxDNA visualization was easily added by following the Three.js WebXR examples. Similarly, it is easy to export the visualized scene to other 3D formats, such as GLTF, for photorealistic rendering (Fig. 1) or 3D printing (Fig. 12).

**FIG. 12.**
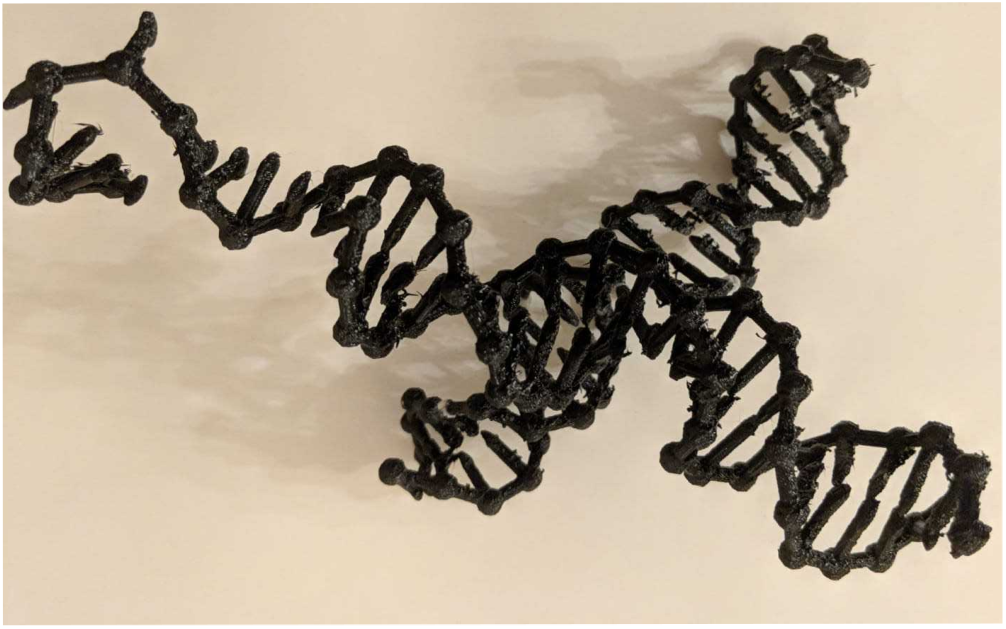
3D printed Holliday junction exported from oxView. OxView supports export to GLTF format that can be opened in a 3D rendering tool Blender and exported to 3D printers or used for creation of more artistic 3D figures of DNA and RNA nanostructures.

The features of oxView and simulation analysis tools are designed to help researchers in DNA and RNA nanotechnology to prototype *in silico* their structures, simplify the design and optimization process, and better understand the functioning of the designed structures. We demonstrated the utility and versatility of the visualization and analysis tools on multiple DNA and RNA nanostructure designs, ranging in size from hundreds to multiple thousands of nucleotides per structure. We also demonstrated that the tools can, in principle, handle structures of sizes over a million nucleotides.

These tools, particularly mean structure calculation and hydrogen bond occupancy, provide significant utility for iterative design of nanostructures. In many structures where unbounded growth is a goal, global curvature of the nanostructure due to subtleties in crossover placement is a significant bottleneck, that is difficult to solve using rational design principles. We have found that the curvature of mean structures calculated from oxDNA simulations (unpublished results) is a good predictor of lattice formation in the laboratory. We also note that mean structures are the best proxy for comparing simulations with cryo-EM structures, which have become important characterizations for 3D nanostructures in the nucleic acid nanotechnology field.

Hydrogen bond occupancy is a good proxy measure for the amount of stress built up in a structure. Even with the speed and level of coarse-graining that oxDNA provides, modelling assembly pathways for large structures remains out of reach for all but the most ambitious simulations^69^. Because of this limitation, we perform simulations with the assumption that the structure forms as designed, and initiate the simulation with all hydrogen bonds present. Designed pairs that become unbonded or find different partners, particularly at junction points, are a good indication for points in the design that are stressed and would benefit from iterative design. In general, we found that successfully published structures had near 100% bond occupancy, while those that were proving difficult to obtain in the lab had regions with low occupancy.

We demonstrated the functionality and versatility of these tools by applying them to a range of DNA and RNA nanostructures, such as DNA and RNA origamis, as well as optimizing and analyzing an RNA tile.

All software discussed here is open-source and freely available through our GitHub under the GNU Public License. Pull requests, bug reports and feature suggestions are welcome, as we hope that these will provide fundamental support long into the future. All tools that were introduced here are documented on their respective GitHub repositories, with examples of use.

## V. DATA AVAILABILITY

The oxDNA simulation code is available online on the oxDNA webpage dna.physics.ox.ac.uk. OxView is available as a web-based application on github.com/sulcgroup/oxdna-viewer. The analysis package can be downloaded from github.com/sulcgroup/oxdna_analysis_tools.

## VI. ACKNOWLEDGEMENTS

We would like to thank R. Hariadi, C. Simmons, and X. Qi for the design files from their previous publications used as examples in this work. We would like to thank members of the Sulc and Yan labs, specifically H. Liu, J. Procyk, and L. Chen for their feedback and beta-testing of the tools presented here.

## VII. FUNDING

This work was supported by the National Science Foundation under grant no. 1931487. JB is supported by the European Unions Horizon 2020 research and innovation program under grant no. 765703.

## REFERENCES

1 N. C. Seeman. Nucleic Acid Junctions and Lattices. Journal of Theoretical Biology, 99:237–247, 1982.

2 H. Yan, T. H. LaBean, L. Feng, and J. H. Reif. Directed nucleation assembly of DNA tile complexes for barcode-patterned lattices. Proceedings of the National Academy of Sciences, 100(14):8103–8108, 2003.

3 T. Gerling, K. F. Wagenbauer, A. M. Neuner, and H. Dietz. Dynamic DNA devices and assemblies formed by shape-complementary, non-base pairing 3D components. Science, 347(6229):1446–1452, 2015.

4 M. Matthies, N. P. Agarwal, E. Poppleton, F. M. Joshi, P. Šulc, and T. L. Schmidt. Triangulated Wireframe Structures Assembled Using Single-Stranded DNA Tiles. ACS Nano, 13(2):1839–1848, 2019.

5 R. Veneziano, S. Ratanalert, K. Zhang, F. Zhang, H. Yan, W. Chiu, and M. Bathe. Designer nanoscale DNA assemblies programmed from the top down. Science, 352(6293):1534–1534, 2016.

6 B. Wei, M. Dai, and P. Yin. Complex shapes self-assembled from single-stranded DNA tiles. Nature, 485(7400):623–626, 2012.

7 D. Han, X. Qi, C. Myhrvold, B. Wang, M. Dai, S. Jiang, M. Bates, Y. Liu, B. An, F. Zhang, et al. Single-stranded DNA and RNA origami. Science, 358(6369):eaao2648, 2017.

8 G. Tikhomirov, P. Petersen, and L. Qian. Fractal assembly of micrometre-scale DNA origami arrays with arbitrary patterns. Nature, 552(7683):67, 2017.

9 A. Gopinath, E. Miyazono, A. Faraon, and P. W. Rothemund. Engineering and mapping nanocavity emission via precision placement of DNA origami. Nature, 535(7612):401, 2016.

10 S. Li, Q. Jiang, S. Liu, Y. Zhang, Y. Tian, C. Song, J. Wang, Y. Zou, G. J. Anderson, J.-Y. Han, et al. A DNA nanorobot functions as a cancer therapeutic in response to a molecular trigger in vivo. Nature biotechnology, 36(3):258, 2018.

11 S. M. Douglas, A. H. Marblestone, S. Teerapittayanon, A. Vazquez, G. M. Church, and W. M. Shih. Rapid prototyping of 3D DNA-origami shapes with caDNAno. Nucleic Acids Research, 37(15):5001–5006, 2009.

12 S. Williams, K. Lund, C. Lin, P. Wonka, S. Lindsay, and H. Yan. Tiamat: A Three-Dimensional Editing Tool for Complex DNA Structures. In A. Goel, F. C. Simmel, and P. Sosík, editors, DNA Computing, pages 90–101, Berlin, Heidelberg, 2009. Springer Berlin Heidelberg.

13 E. Benson, A. Mohammed, J. Gardell, S. Masich, E. Czeizler, P. Orponen, and B. Högberg. DNA rendering of polyhedral meshes at the nanoscale. Nature, 523(7561):441–444, 2015.

14 E. Benson, A. Mohammed, A. Bosco, A. I. Teixeira, P. Orponen, and B. Hogberg. Computer-Aided Production of Scaffolded DNA Nanostructures from Flat Sheet Meshes. Angewandte Chemie International Edition, 55(31):8869–8872, 2016.

15 E. de Llano, H. Miao, Y. Ahmadi, A. J. Wilson, M. Beeby, I. Viola, and I. Barisic. Adenita: Interactive 3D modeling and visualization of DNA Nanostructures. bioRxiv, page 849976, 2019.

16 C.-M. Huang, A. Kucinic, J. Johnson, H.-J. Su, and C. E. Castro. Hybrid top-down and bottom-up approach for engineering DNA assemblies. in preparation.

17 R. Veneziano, S. Ratanalert, K. Zhang, F. Zhang, H. Yan, W. Chiu, and M. Bathe. Designer nanoscale DNA assemblies programmed from the top down. Science, 352(6293):1534, 2016.

18 H. Jun, F. Zhang, T. Shepherd, S. Ratanalert, X. Qi, H. Yan, and M. Bathe. Autonomously designed free-form 2D DNA origami. Science advances, 5(1):eaav0655, 2019.

19 A. Suma, A. Stopar, A. W. Nicholson, M. Castronovo, and V. Carnevale. Allosteric modulation of local reactivity in DNA origami. BioRxiv, page 640847, 2019.

20 C. Maffeo, J. Yoo, and A. Aksimentiev. De novo reconstruction of DNA origami structures through atomistic molecular dynamics simulation. Nucleic acids research, 44(7):3013–3019, 2016.

21 P. Fonseca, F. Romano, J. S. Schreck, T. E. Ouldridge, J. P. Doye, and A. A. Louis. Multi-scale coarse-graining for the study of assembly pathways in DNA-brick self-assembly. Journal of Chemical Physics, 148(13), 2018.

22 A. Reinhardt and D. Frenkel. Numerical evidence for nucleated self-assembly of DNA brick structures. Physical review letters, 112(23):238103, 2014.

23 B. E. Snodin, J. S. Schreck, F. Romano, A. A. Louis, and J. P. Doye. Coarse-grained modelling of the structural properties of DNA origami. Nucleic Acids Research, 47(3):1585–1597, 2019.

24 T. E. Ouldridge, P. Šulc, F. Romano, J. P. Doye, and A. A. Louis. DNA hybridization kinetics: Zippering, internal displacement and sequence dependence. Nucleic Acids Research, 41(19):8886–8895, 2013.

25 J. S. Schreck, T. E. Ouldridge, F. Romano, P. Šulc, L. P. Shaw, A. A. Louis, and J. P. K. Doye. DNA hairpins destabilize duplexes primarily by promoting melting rather than by inhibiting hybridization. Nucleic Acids Research, 43(13):6181–6190, 2015.

26 P. Šulc, T. E. Ouldridge, F. Romano, J. P. Doye, and A. A. Louis. Simulating a burnt-bridges DNA motor with a coarse-grained DNA model. Natural Computing, 13(4):535–547, 2014.

27 T. E. Ouldridge, R. L. Hoare, A. A. Louis, J. P. Doye, J. Bath, and A. J. Turberfield. Optimizing DNA nanotechnology through coarse-grained modeling: a two-footed DNA walker. ACS nano, 7(3):2479–2490, 2013.

28 T. E. Ouldridge, A. A. Louis, and J. P. Doye. Structural, mechanical, and thermodynamic properties of a coarse-grained DNA model. The Journal of chemical physics, 134(8):02B627, 2011.

29 B. E. Snodin, F. Randisi, M. Mosayebi, P. Šulc, J. S. Schreck, F. Romano, T. E. Ouldridge, R. Tsukanov, E. Nir, A. A. Louis, et al. Introducing improved structural properties and salt dependence into a coarse-grained model of DNA. The Journal of chemical physics, 142(23):06B613_1, 2015.

30 P. Šulc, F. Romano, T. E. Ouldridge, L. Rovigatti, J. P. Doye, and A. A. Louis. Sequence-dependent thermodynamics of a coarse-grained DNA model. The Journal of chemical physics, 137(13):135101, 2012.

31 P. Šulc, F. Romano, T. E. Ouldridge, J. P. Doye, and A. A. Louis. A nucleotide-level coarse-grained model of RNA. Journal of Chemical Physics, 140(23), 2014.

32 J. P. Doye, T. E. Ouldridge, A. A. Louis, F. Romano, P. Sulc, C. Matek, B. E. K. Snodin, L. Rovigatti, J. S. Schreck, R. M. Harrison, and W. P. J. Smith. Coarse-graining DNA for simulations of DNA nanotechnology. Physical Chemistry Chemical Physics, 15(47):20395–20414, 2013.

33 C. Maffeo and A. Aksimentiev. MrDNA: A multi-resolution model for predicting the structure and dynamics of nanoscale DNA objects. bioRxiv, page 865733, 2019.

34 D.-N. Kim, F. Kilchherr, H. Dietz, and M. Bathe. Quantitative prediction of 3D solution shape and flexibility of nucleic acid nanostructures. Nucleic acids research, 40(7):2862–2868, 2011.

35 C. E. Castro, F. Kilchherr, D.-N. Kim, E. L. Shiao, T. Wauer, P. Wortmann, M. Bathe, and H. Dietz. A primer to scaffolded DNA origami. Nature methods, 8(3):221, 2011.

36 D. M. Hinckley, G. S. Freeman, J. K. Whitmer, and J. J. De Pablo. An experimentally-informed coarse-grained 3-site-per-nucleotide model of DNA: Structure, thermodynamics, and dynamics of hybridization. Journal of Chemical Physics, 139(14):1–16, 2013.

37 M. C. Engel, D. M. Smith, M. A. Jobst, M. Sajfutdinow, T. Liedl, F. Romano, L. Rovigatti, A. A. Louis, and J. P. Doye. Force-induced unravelling of DNA origami. ACS nano, 12(7):6734–6747, 2018.

38 R. Sharma, J. S. Schreck, F. Romano, A. A. Louis, and J. P. Doye. Characterizing the motion of jointed DNA nanostructures using a coarse-grained model. ACS nano, 11(12):12426–12435, 2017.

39 F. Hong, S. Jiang, X. Lan, R. P. Narayanan, P. Šulc, F. Zhang, Y. Liu, and H. Yan. Layered-crossover tiles with precisely tunable angles for 2D and 3D DNA crystal engineering. Journal of the American Chemical Society, 140(44):14670–14676, 2018.

40 C. Matek, P. Šulc, F. Randisi, J. P. Doye, and A. A. Louis. Coarse-grained modelling of supercoiled RNA. The Journal of chemical physics, 143(24):243122, 2015.

41 T. E. Ouldridge, P. Šulc, F. Romano, J. P. Doye, and A. A. Louis. DNA hybridization kinetics: zippering, internal displacement and sequence dependence. Nucleic acids research, 41(19):8886–8895, 2013.

42 A. Suma, E. Poppleton, M. Matthies, P. Šulc, F. Romano, A. A. Louis, J. P. Doye, C. Micheletti, and L. Rovigatti. TacoxDNA: A user-friendly web server for simulations of complex DNA structures, from single strands to origami. Journal of computational chemistry, 40(29):2586–2595, 2019.

43 S. Van Der Walt, S. C. Colbert, and G. Varoquaux. The NumPy array: A structure for efficient numerical computation. Computing in Science and Engineering, 13(2):22–30, 2011.

44 J. D. Hunter. Matplotlib: A 2D graphics environment. Computing in Science & Engineering, 9(3):90–95, 2007.

45 P. J. Cock, T. Antao, J. T. Chang, B. A. Chapman, C. J. Cox, A. Dalke, I. Friedberg, T. Hamelryck, F. Kauff, B. Wilczynski, and M. J. De Hoon. Biopython: Freely available Python tools for computational molecular biology and bioinformatics. Bioinformatics, 25(11):1422–1423, 2009.

46 F. Pedregosa, R. Weiss, and M. Brucher. Scikit-learn: Machine Learning in Python. Journal of Machine Learning Research, 12:28252830, 2011.

47 M. M. McKerns, L. Strand, T. Sullivan, A. Fang, and M. A. G. Aivazis. Building a Framework for Predictive Science. Proceedings of the 10th Python in Science Conference, pages 1–12, 2011.

48 B. E. Snodin, F. Randisi, M. Mosayebi, P. Šulc, J. S. Schreck, F. Romano, T. E. Ouldridge, R. Tsukanov, E. Nir, A. A. Louis, and J. P. Doye. Introducing improved structural properties and salt dependence into a coarse-grained model of DNA. Journal of Chemical Physics, 142(23), 2015.

49 L. Rovigatti, P. Šulc, I. Z. Reguly, and F. Romano. A comparison between parallelization approaches in molecular dynamics simulations on GPUs. Journal of computational chemistry, 36(1):1–8, 2015.

50 B. E. Snodin, F. Romano, L. Rovigatti, T. E. Ouldridge, A. A. Louis, and J. P. Doye. Direct simulation of the self-assembly of a small DNA origami. ACS nano, 10(2):1724–1737, 2016.

51 J. Russo, P. Tartaglia, and F. Sciortino. Reversible gels of patchy particles: role of the valence. The Journal of chemical physics, 131(1):014504, 2009.

52 S. Whitelam and P. L. Geissler. Avoiding unphysical kinetic traps in Monte Carlo simulations of strongly attractive particles. Journal of Chemical Physics, 127(15), 2007.

53 M. Ester, H.-P. Kriegel, J. Sander, X. Xu, et al. A density-based algorithm for discovering clusters in large spatial databases with noise. In Kdd, volume 96, pages 226–231, 1996.

54 M. Schickinger, M. Zacharias, and H. Dietz. Tethered multifluorophore motion reveals equilibrium transition kinetics of single DNA double helices. Proceedings of the National Academy of Sciences, 115(39):201800585, 2018.

55 C.-M. Huang, A. Kucinic, J. V. Le, C. E. Castro, and H.-J. Su. Uncertainty quantification of a DNA origami mechanism using a coarse-grained model and kinematic variance analysis. Nanoscale, 11(4):1647–1660, 2019.

56 D. Baraff. An introduction to physically based modeling: Rigid Body Simulation I Unconstrained Rigid Body Dynamics, 1997.

57 S. M. Douglas, H. Dietz, T. Liedl, B. Högberg, F. Graf, and W. M. Shih. Self-assembly of DNA into nanoscale three-dimensional shapes. Nature, 459(7245):414, 2009.

58 H. J. Berendsen, D. van der Spoel, and R. van Drunen. GROMACS: A message-passing parallel molecular dynamics implementation. Computer Physics Communications, 91(1-3):43–56, 1995.

59 W. Kabsch. A solution for the best rotation to relate two sets of vectors. crystallographica Section A: Crystal Physics, Diffraction, Theoretical and General Crystallography, 32:922–923, 1976.

60 G. Young and A. S. Householder. Discussion of a set of points in terms of their mutual distances. Psychometrika, 3(1):19–22, 1938.

61 I. Borg and P. Groenen. Modern multidimensional scaling: Theory and applications. Journal of Educational Measurement, 40(3):277–280, 2003.

62 H. Dietz, S. M. Douglas, and W. M. Shih. Folding DNA into Twisted and Curved Nanoscale Shapes Hendrik. Science, 325(5941):725–730, 2009.

63 M. A. Baker, A. J. Tuckwell, J. F. Berengut, J. Bath, F. Benn, A. P. Duff, A. E. Whitten, K. E. Dunn, R. M. Hynson, A. J. Turberfield, and L. K. Lee. Dimensions and Global Twist of Single-Layer DNA Origami Measured by Small-Angle X-ray Scattering. ACS Nano, 12(6):5791–5799, 2018.

64 E. F. Pettersen, T. D. Goddard, C. C. Huang, G. S. Couch, D. M. Greenblatt, E. C. Meng, and T. E. Ferrin. UCSF Chimera - A visualization system for exploratory research and analysis. Journal of Computational Chemistry, 25(13):1605–1612, 2004.

65 D. Han, X. Qi, C. Myhrvold, B. Wang, M. Dai, S. Jiang, M. Bates, Y. Liu, B. An, F. Zhang, H. Yan, and P. Yin. Single-stranded DNA and RNA origami. Science, 358(6369), 2017.

66 V. Schomaker, J. Waser, R. E. Marsh, and G. Bergman. To fit a plane or a line to a set of points by least squares. Acta Crystallographica, 12(8):600–604, 1959.

67 C. C. David and D. J. Jacobs. Principal Component Analysis: A Method for Determining the Essential Dynamics of Proteins. Methods in Molecular Biology, 1084:193–226, 2014.

68 X. Qi, F. Zhang, Z. Su, S. Jiang, D. Han, B. Ding, Y. Liu, W. Chiu, P. Yin, and H. Yan. Programming molecular topologies from single-stranded nucleic acids. Nature communications, 9(1):1–9, 2018.

69 B. E. Snodin, F. Romano, L. Rovigatti, T. E. Ouldridge, A. A. Louis, and J. P. Doye. Direct Simulation of the Self-Assembly of a Small DNA Origami. ACS Nano, 10(2):1724–1737, 2016.

